# Identification of cross-reacting IgG hotspots to prevent immune evasion of SARS-CoV-2 variants

**DOI:** 10.1101/2023.06.27.546805

**Authors:** Marek Harhala, Katarzyna Gembara, Krzysztof Baniecki, Aleksandra Pikies, Artur Nahorecki, Natalia Jędruchniewicz, Zuzanna Kaźmierczak, Izabela Rybicka, Tomasz Klimek, Wojciech Witkiewicz, Kamil Barczyk, Marlena Kłak, Krystyna Dąbrowska

## Abstract

The major factor that shapes the global perspective for increase or diminution of successive pandemic waves of COVID-19 is the immunological protection. The SARS-CoV-2 virus constantly develops new variants, and capability of immune evasion is among the major factors that promote variant spreading in the human population. After two years of the pandemic and virus evolution, it is almost impossible to explain effects of all possible combinations different viral strains, a few types of vaccinations or new variants infecting an individual patient. Instead of variant-to-variant comparisons, identification of key protein regions linked to immune evasion could be efficient.

Here we report an approach for experimental identification of SARS-CoV-2 protein regions that (i) have characteristics of cross-reacting IgG hot-spots, and (ii) are highly immunogenic. Cross-reacting IgG hot spots are regions of protein frequently recognized in many variants by cross-reacting antibodies. Immunogenic regions efficiently induce specific IgG production in SARS-CoV-2 infected patients. We determined four regions that demonstrate both significant immunogenicity and the activity of a cross-reacting IgG hot-spot in protein S, and two such regions in protein N. Their distribution within the proteins suggests that they may be useful in vaccine design and in serological diagnostics of COVID-19.

## Introduction

Efficient immunological protection in COVID-19 is currently the major factor that shapes the global perspective for further pandemic waves growing or receding. The SARS-CoV-2 virus has constantly been producing a multitude of variants; many of these variants, such as Delta or Omicron, induced pandemic waves obviously using immune evasion to spread within the global population ^1,2^. Importantly, antigenic drift that is driven by the pressure from specific immunity that is exerted on the virus markedly contributes to selection of new variants able to evade immunological control ^3^. For these reasons, the hope for efficient control and suppression of the pandemic simply due to global vaccinations and immunological memory of previous infections has been tempered.

Virus-specific antibodies are in fact a pool of antibodies targeting different structural proteins of the virus. Structural proteins of SARS-CoV-2 include nucleocapsid protein (protein N), spike protein (protein S), envelope protein (protein E), and membrane protein (protein M) ^4^. The most important target for the immune response is protein S, due to the virus-blocking potential of antibodies targeting this protein. This protein mediates binding to the ACE2 receptor; thus antibodies targeting S protein may neutralize the virus. S protein is relatively large, 1273 amino acids (aa) long (acc. no.: YP_009724390.1) and it consists of two major domains: S1 and S2. The S1 subunit function is binding to receptors on human cells. Its major parts are: the N-terminal domain (NTD) and receptor binding domain (RBD). The S2 subunit, in turn, is engaged in the viral and host cell membranes’ fusion. S2 consists of: the cytoplasmic tail (CT), transmembrane domain (TM), heptad repeat 1 (HR1), heptad repeat 2 (HR2), fusion peptide (FP), central helix (CH), and connector domain (CD). Also, cleavage site S1/S2 plays the key role in virus entry to a human host cell. Linear epitopes on S protein have been demonstrated to elicit neutralizing antibodies in COVID-19 patients ^5^. N protein, in turn, seems to be a key target for serological diagnostics. This relatively conserved protein has commonly been used in serological diagnostics, particularly to confirm SARS-CoV-2-related etiology of already cleared infections, since N protein is unique for SARS-CoV-2, highly immunogenic, and not included in anti-COVID-19 vaccines ^6^. Two other proteins, M and E, seem to have lower significance as potential targets for specific antibodies. However, one cannot exclude a role of these antibodies in virus inhibition, for instance via complement system activation or facilitating phagocytosis.

After two years of the pandemic and virus evolution within the human population, it is almost impossible to explain all possible combinations of previous infections with different variants, a few types of vaccinations, and new variants that may infect an individual patient. These combinations further generate myriads of possibilities at the population level. Thus, instead of variant-to-variant comparisons, identification of key protein regions that are linked to immune evasion could be more helpful for predictions on pandemic potential in new SARS-CoV-2 variants. Here we report these experimentally identified regions (linear epitopes) in SARS-CoV-2 structural proteins, including protein S as the major target for immunological protection and protein N widely used for serological diagnostics. The *cross-reacting IgG hot-spots* have been identified by a high-throughput assay of cross-reactivity of antibodies elicited by SARS-CoV-2, based on the modified VirScan technology ^7^. This approach employs a phage display library of epitopes and it allows for significant extension of the most popular method for B-cell epitope identification (by microarrays) and for massive testing of multitude epitopes; here, 96 623 oligopeptides derived from available variants of SARS-CoV-2 have been tested (Supplementary Data 1).

## Materials and methods

### Serum donors

Blood serum was collected from patients hospitalized due to COVID-19: PCR confirmed infection with SARS-CoV-2 (at least 1 month after infection), non-vaccinated, over 18 years old, hospitalized in COVID-19 ward in Healthcare Centre Bolesławiec, Poland (alpha variant: n=15, delta variant: n=8).

### Bioethics statements

The study was conducted in accordance with the principles of the Declaration of Helsinki. The research was approved by the local Bioethical Commission of the Regional Specialist Hospital in Wroclaw (approval number: KB/02/2020, policy No. COR193657). During the individual interview, all information about the study was provided and written consent was obtained from each participant. The written consent was accepted by the local Bioethical Commission of the Regional Specialist Hospital in Wroclaw (approval number: KB/02/2020).

### Blood samples

Blood samples were collected in test tubes (BD SST II Advance), left to clot for 1 hour at room temperature (RT), and separated from the clot by centrifugation (15 min, 2000 g, RT) and then stored at – 20°C for further use.

### Identification of SARS-CoV-2 variants infecting investigated patients

In selected hospitalized patients, SARS-CoV-2 variants were identified by the targeted sequencing approach with the Ion AmpliSeq SARS-CoV-2 Insight Research Assay (Thermo Fisher Scientific, Europe, according to the manufacturer’s instructions). Briefly, patients were sampled by nasopharyngeal swabbing, viral RNA was isolated on silica spin columns (QIAquick) and the viral load was quantified by the qPCR standard diagnostic test. Ct values were used to design library preparation parameters, which included standard retrotranscription. Library preparation and sequencing chip loading were completed on Ion Chef. Sequencing was completed on Ion GeneStudio S5, and analyzed with the software provided by the manufacturer. Sequences of sufficient quality were uploaded to the public database GISAID (https://www.gisaid.org/).

### Epitope analysis

Identification of oligopeptides interacting with specific IgGs was performed in accordance with the modified protocol published by Xu et al.^7^ and adapted for our research using coding sequences of the investigated SARS-CoV-2 variants as the source for library design ^7^. Part of a library representing protein variants was created from proteomes of SARS-CoV-2 variants downloaded on 07.04.2021 from the Identical Protein Groups Database, National Center for Biotechnology Information. Another part of a library representing the reference proteome of the SARS-CoV-2 virus (prot. ID: UP000464024) was created from the Proteome Database at UniProt Proteome resources and set of oligopeptides was created by single and triple alanine substitution, similarly to original procedure ^7^.

Variants of each investigated reference protein were aligned with Clustal Omega software ^8–10^. Each such alignment and protein library consisting of reference proteomes was virtually cut into 56 aa long fragments tailing through protein sequences, starting every 28 aa from the first amino acid. The peptide library, consisting of protein variants in Identical Protein Group, was cleared from all incomplete sequences (containing missing fragments or undetermined amino acids). If a specific protein showed a gap in the alignment, oligopeptides covering this place were shorter than 56 aa. The start position of each oligopeptide in the reference proteins N, S, M, and E of SARS-CoV-2 is presented in Supplementary Table S1-S5.

Sequences of all oligopeptides were reverse-translated into DNA sequences using codons optimized for expression in *E. coli*. The oligopeptide library was synthesized using the SurePrint technology for nucleotide printing (Agilent). These oligonucleotides were used to create a phage display library using the T7Select 415-1 Cloning Kit (Merck Millipore). Immunoprecipitation of the library was performed in accordance with a previously published protocol^7,11^. Briefly, the phage library was amplified in a standard culture as described in the manufacturer’s manual, then purified by hollow fiber dialysis against Phage Extraction Buffer (20 mM Tris-HCl, 100 mM NaCl, 6 mM MgSO4, pH 8.0). All plastic containers (96-well plates) used for immunoprecipitation were prepared by blocking with 3% Bovine Serum Albumin (BSA) in TBST buffer overnight on a rotator (50 rpm, 4°C). A sample representing an average of 10^5^ copies of each clone in 250 µL was mixed with 1 µL of human serum (two technical replicates were applied) and incubated overnight at 4°C with rotation (50 rpm). A 20 μL aliquot of a 1:1 mixture of Protein A and Protein G Dynabeads (Invitrogen) was added and incubated for 4 h at 4°C with rotation (50 rpm). Liquid in all wells was separated from Dynabeads on a magnetic stand and removed. Beads were washed 5 times with 280 µL of a wash buffer (50 mM Tris-HCl, pH 7.5, 150 mM NaCl, 0.1% Tween-20) and beads were resuspended in 60 µL of water to elute the immunoprecipitated bacteriophages from the beads.

The immunoprecipitated part of the library (for each patient-derived sample), as well as 20 samples representing the library before immunoprecipitation (*input samples*), was then used for amplification of the insert region according to the manufacturer’s instructions with a Phusion Blood Direct PCR Kit (Thermo Fisher Scientific). A second round of PCR was carried out with the IDT for Illumina UD indexes (Illumina Corp.) to add adapter tags. Sequencing of the amplicons was completed using Illumina next generation sequencing (NGS) technology (Genomed, Warszawa). A full list of oligopeptides in the tested library (cloned) is given in Supplementary Data 1.

### Sequencing data analysis

Sequenced amplicons were mapped to the original nucleotide library sequences by the bowtie2 software similarly as described by Xu et al. ^7,12^. NGS sequencing reads (after removal of sequences added in PCR) were mapped to the full list of oligonucleotides as designed for the library synthesis as indexes (options: end-to-end mode, ‘-q -5 9 --no-unal --no-hd --no-sq --ignore-quals --mp 3 --rdg 150,100 --rfg 150,100 - -score-min L,-0.6,-0.6’) ^12,13^. The number of hits that mapped to each reference sequence (only the highest score for each read) was counted (*count, c*).

The signal in each sample was calculated/normalized according to the formula (1):

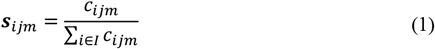

s – signal of i-th sequence in j-th serum sample and m-th technical replicate

c – count, a number of reads mapped to i-th sequence in m-th technical replicate of j-th serum sample

I – set of all reference sequences (used as indexes in mapping by the bowtie2 software)

Signals in each *input sample* used as negative control samples (n=20, amplified and purified phage library sequenced before immunoprecipitation) were calculated. In *input samples*, the average log10 of a signal and its standard deviation (SD) for each tested sequence were used as *control signal* for later calculation of p-values. As for tested samples, the log10(signal) was determined for each detected sequence (*count*>0). In order to avoid false positive results we set the signal of undetected oligonucleotides (*s*) in a tested sample to the minimum measured for this sample value and its standard deviation to the average value of SD in *input samples*.

Each log10(signal) in tested samples was compared to the *control signal* of the same sequence in *input samples* and the p-value was calculated (assuming normal distribution). Only signals detected with a non-zero count and p-value<0.05 in both technical replicates of a sample were recognized as *positive* (significantly enriched). *Relative signal* (enrichment): an average signal in technical replicates of a sample divided by the average signal in ‘input samples’ of the same sequence resulted in a signal ratio (*relative signal*). The above procedure was tested on *input samples* as controls and proved to yield less than a 5% chance for a false *positive* (significantly enriched) sequence in each sample.

### Immunogenicity and cross-reacting IgG hot-spot determination – statistical model

Determination of immunogenicity and cross-reacting IgG hot-spot was done by adopting binomial distribution. This distribution (provided with *chance* and size) allows one to calculate the chance of randomly observing a specific amount of positive outcomes in a series of trials. The assumption is that every patient has the same chance of recognizing each measured oligopeptide and its variant separately.

For each group (Alpha/Delta, N/S-protein) in both experiments (immunogenicity research and hot-spot analysis) we used the same protocol separately. First, we calculated *a chance* by dividing the sum of all detected (positive) oligopeptides in each patient in a group by all tested patients and all tested oligopeptides/variants. A *size* is the amount of patients in a researched group (immunogenicity) or the number of tested variants of a fragment in question (cross-reacting hot-spots). Then in R language we calculated the chance for a random occurrence of an observed number of positive results for each tested protein fragment. Fragments whose chance for a random occurrence of a measured result is below 0.05 were indicated in the figures with one star and results below 0.001 were indicated with double stars. Exact p-values are provided (Supplementary Table S1-S5) along with graphical comparison of the statistical model used and observed data (Supplementary Figure S1).

## Results

Immunoreactivity of specified regions (oligopeptides) within SARS-CoV-2 proteins was assessed by the modified VirScan technology, presented on Figure 1 ^7^. A phage display library derived from fragments of N, S, M, E proteins and variants of these proteins (downloaded from National Center for Biotechnology Information on 07.04.2021) was constructed and used for immunoprecipitation with sera from patients hospitalized due to COVID-19 (SARS-CoV-2 variants causing those infections were identified; they all belonged to Alpha or Delta type). After NGS of the immunoprecipitated libraries, two types of analysis were conducted: (i) identification of regions frequently recognized in many variants by cross-reacting antibodies (*IgG hot-spots*), (ii) identification of immunogenic regions within structural proteins of SARS-Cov-2.

**Figure 1:**
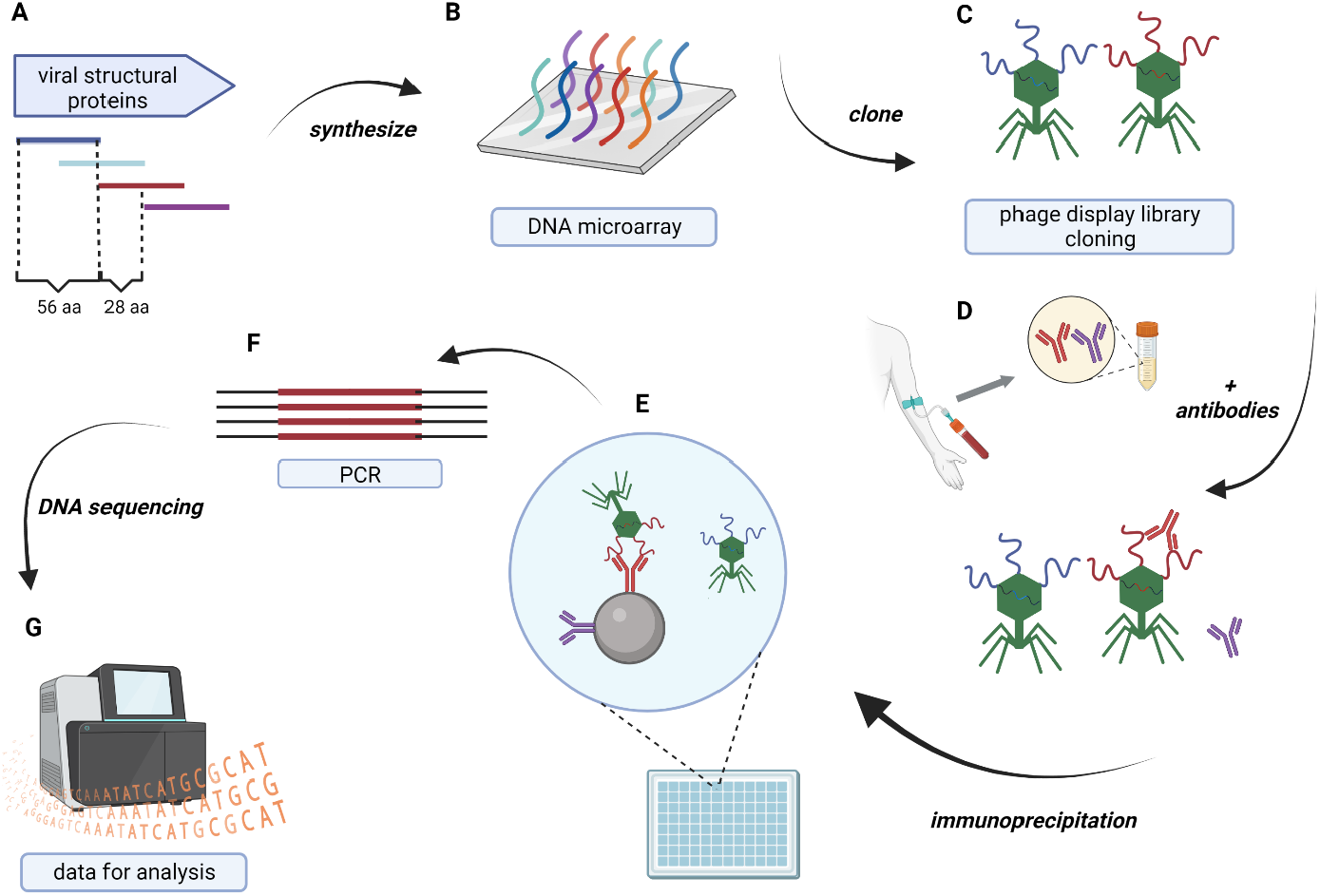
VirScan technology used for immunoreactivity measurements. (a) In silico design of oligopeptides representing chosen proteomes, (b) synthesis of oligonucleotides coding for the peptides,(c) constructing of phage display library of SARS-CoV-2-derived peptides, (d) reaction of the library with patients sera, (e) immunoprecipitation with magnetic beads binding Fc fragments of antibodies, (f) amplification by PCR, (g) NGS sequencing, modified from Xu et al.^7^.

### Some protein regions can be altered without affecting their recognition by patients IgG antibodies

Protein regions frequently recognized by cross-reacting IgG antibodies in many SARS-CoV-2 variants in spite of amino acid differences were identified as *cross-reacting IgG hot-spots* (Figure 2, Supplementary Tables S1 and S2). The library of oligopeptides representing structural SARS-CoV-2 proteins of multiple variants (96 623 oligopeptides, including Omicron-derived sequences, Supplementary Data 1) was immunoprecipitated with sera from the investigated patients. The fraction of variants recognized by patients’ IgG was calculated (Figure 2) and normalized to a total number each oligopeptide variants cloned into the library (this number differed between protein regions; see Table S1 and S2). A high fraction of cross-reacting variants (over 20%, p<0.05) indicates a *hot-spot*, while low scores indicate regions where new variants efficiently escaped antibodies induced by Alpha or Delta. The majority of cross-reacting *IgG hot-spots* were identified in proteins N and S (Figure 2), while only one was identified in protein M and none in protein E (Supplementary Table S3). High similarity can be observed between the *IgG hot-spots* identified by Alpha or Delta-induced IgG, thus suggesting regularities in the immune evasion of the virus.

**Figure 2.**
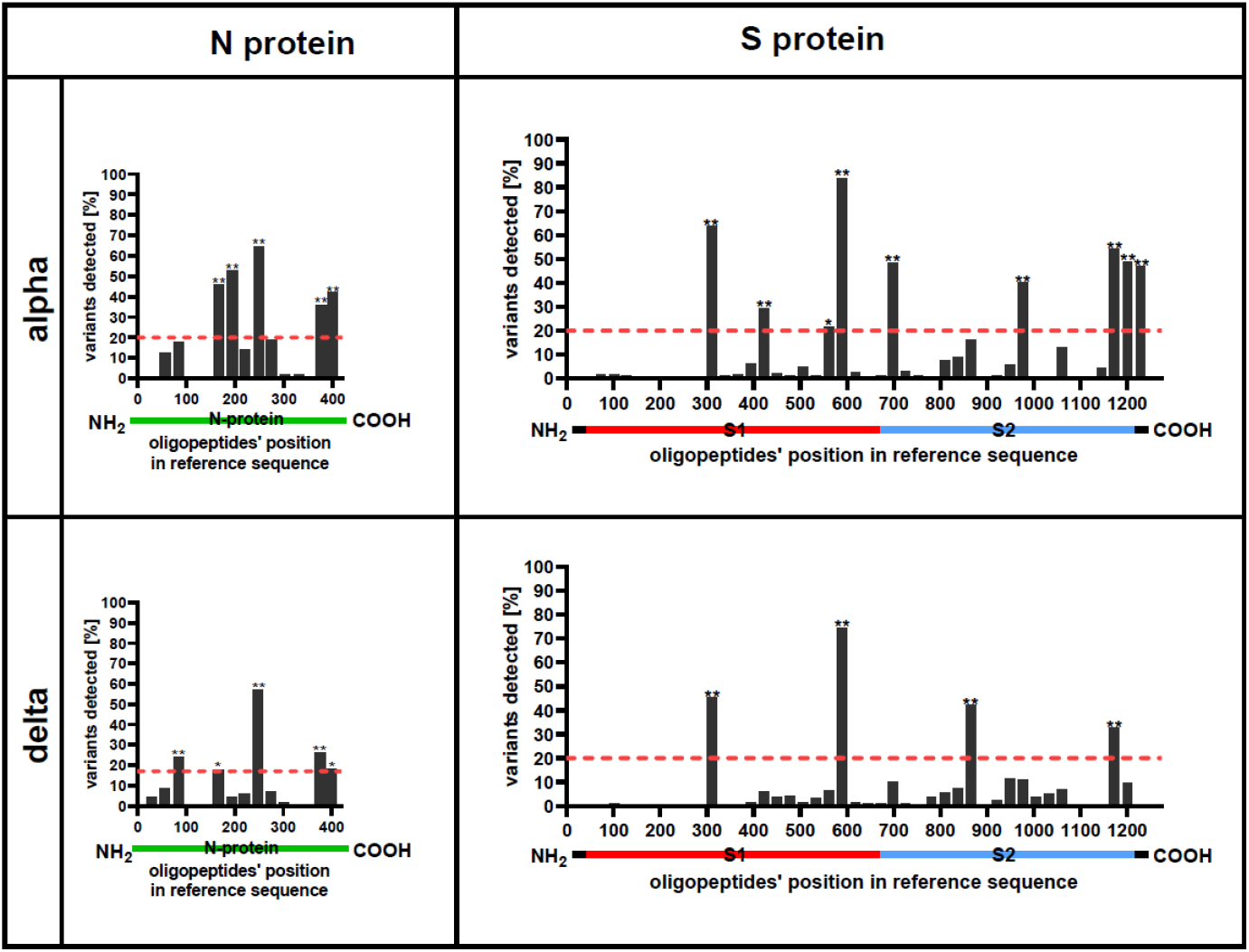
Cross-reacting *IgG hot-spots* in N and S protein of SARS-CoV-2. Cross-reacting IgG hot spots are regions of protein frequently recognized in many variants by cross-reacting antibodies. Cross-reactions of IgG antibodies were identified by immunoprecipitation of the library containing SARS-CoV-2 variants’ oligopeptides (VirScan technology). Immunoprecipitation was conducted with sera from patients hospitalized due to SARS-CoV-2 infection (Alpha or Delta). X-axis represents sequentially arranged oligopeptides that sum up to the whole protein sequence, presented from N-terminus to C-terminus. Each bar represents the fraction of tested viral variants (see Table S4 and S5) that were recognized by antibodies in patients’ sera (N_alpha_ = 15, N_delta_=8). Green lines represent N protein. Red, blue and black lines under the plots represent S protein. * - p<0.05, ** - p<0.001, one-tail. Red dotted line indicates cut-off p<0.05. P-values are calculated between experimental data and statistical binomial distribution model assuming random distribution of hot-spot regions in proteins.

### Immunogenicity differs between protein regions

Further, identification of the immunogenic regions was conducted by detecting homogenic IgG-oligopeptide reactions. These reactions represented binding between oligopeptides derived from reference SARS-CoV-2 virus and antibodies induced in patients hospitalized due to COVID-19. Fractions of patients whose sera contained IgG targeting defined regions of the proteins N and S are presented in Figure 3. In the majority of cases, immunogenic regions are similar between tested SARS-CoV-2 variants (Figure 3, Supplementary Table S4 and Supplementary Table S5). In proteins M and E, in turn, regions of significant immunogenicity were not detected (Table S3).

**Figure 3.**
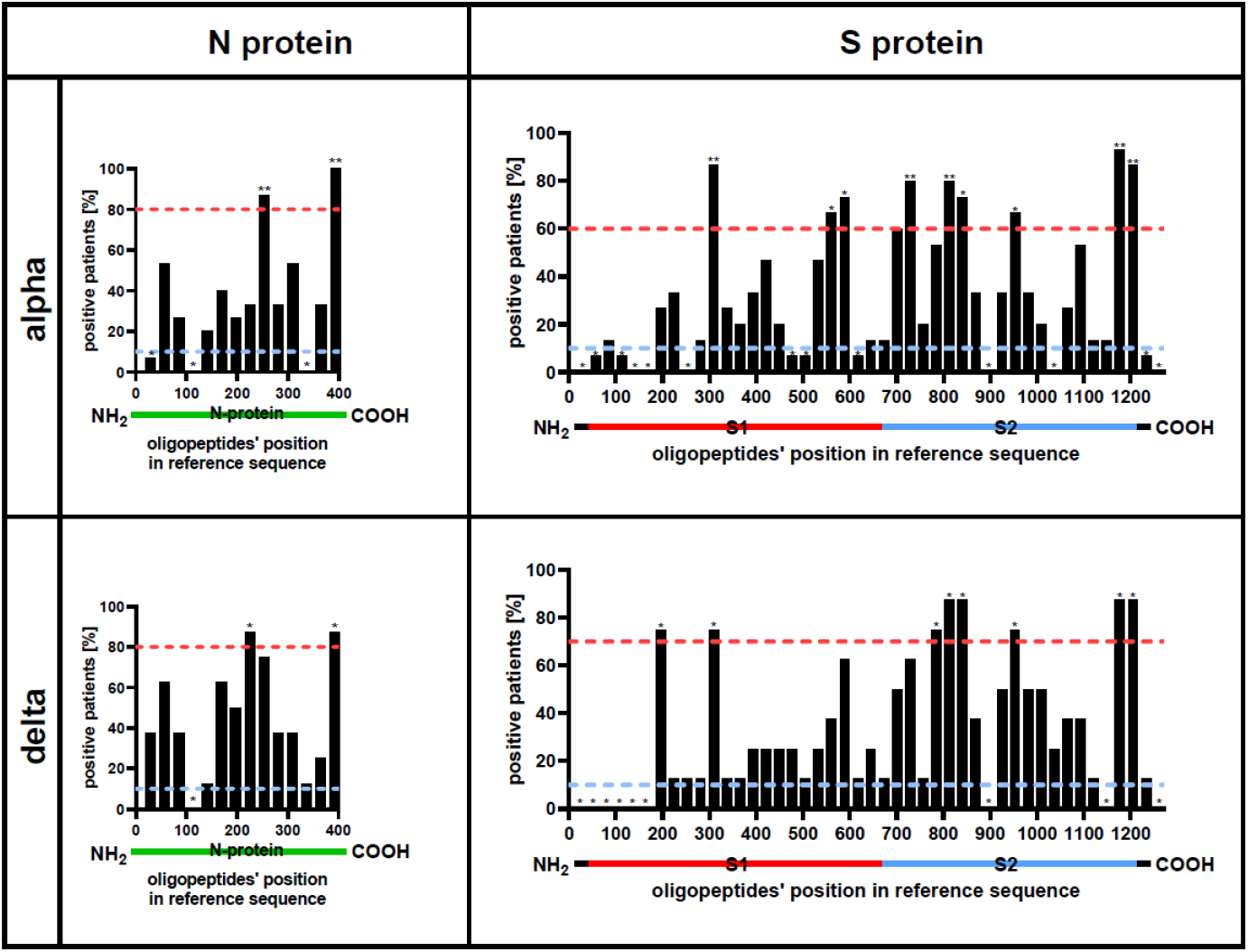
Immunogenic regions of SARS-CoV-2 protein N and S. Immunogenic regions efficiently induce specific IgG production in SARS-CoV-2 infected patients. Immunogenic regions were identified by immunoprecipitation of oligopeptide library representing Alpha and Delta SARS-CoV-2 variants (VirScan technology). Immunoprecipitation was conducted with sera from patients hospitalized due to SARS-CoV-2 infection (Alpha or Delta). X-axis represents sequentially arranged oligopeptides that sum up to the whole protein sequence, presented from N-terminus to C-terminus. Each bar represents the fraction of tested patients who were positive for antibodies specific for a relevant region within SARS-CoV-2 proteins (N_alpha_ = 15, N_delta_=8). Green line under the plot represents N protein. Red, blue and black lines under the plot represent S protein. * - p<0.05, ** - p<0.001. Red and blue dotted lines indicate upper and lower cut-off p<0.05 (two-tail), respectively. P-values are calculated between experimental data and statistical binomial distribution model assuming random distribution of hot-spot regions in proteins.

### Some protein regions show both high immunogenicity and presence of cross-reactive hot-spots

Immunogenic regions (Figure 3, Table S4 and S5) are not fully consistent with cross-reactivity *IgG hot-spot* regions (Figure 2, Tables S1 and S2). In protein N, there are two regions that demonstrate both significant immunogenicity and the activity of a cross-reacting *IgG hot-spot* located between aa positions 197 and 280 or between 358 and 419. In protein S, four regions that demonstrate both significant immunogenicity and the activity of a cross-reacting *IgG hot-spot* were identified: between 281 and 337, 533 and 617, 925 and 1004, and 1145 and 1232. Thus, immunogenic regions with *IgG hot-spot* properties are located both in S1 and S2 domains of protein S. The first and the second are located at the C-terminus of the NTD domain and in the RBD domain, respectively. Their comparison and the list is presented (Figure 4, Table 1). This strongly suggests that they can be useful as virus-neutralizing antibody targets, thus potentially being important in vaccine composition or in monoclonal antibody production. This is due to the increased efficacy of these regions in antibody induction (immunogenicity) and at the same time decreased probability of natural immune evasion demonstrated by multiple variants of the SARS-CoV-2 virus. The last two immunogenic regions with *IgG hot-spot* properties are located within HR1 and HR2 domains, which have also been proposed as efficient targets for antibodies neutralizing the virus ^14^.

**Table 1.** Highly immunogenic regions with strong cross-reacting *IgG hot-spot* in proteins N and S. Immunogenic regions efficiently induce specific IgG production in SARS-CoV-2 infected patients. Immunogenic regions were identified by immunoprecipitation of the oligopeptide library representing Alpha and Delta SARS-CoV-2 variants (VirScan technology). Cross-reacting IgG hot spots are regions of protein frequently recognized in many variants by cross-reacting antibodies. IgG antibodies were identified by immunoprecipitation of the library containing SARS-CoV-2 variants’ oligopeptides (VirScan technology). Immunoprecipitation was conducted with sera from patients hospitalized due to SARS-CoV-2 infection (Alpha or Delta). Listed below are regions of high or very high immunogenicity (p<0.05 and p<0.001, respectively) and strong cross-reacting hot-spots (p<0.001). P-value calculated between experimental data and statistical binomial distribution model assuming random distribution of immunogenicity and hot-spot regions in proteins.

|  | reference protein fragment<br>[amino acid position] |  | Immunogenicity |  | Cross-reacting IgG hot spots |  |
| --- | --- | --- | --- | --- | --- | --- |
|  | start | end | alpha | delta | alpha | delta |
| **N-protein** | 197 | 280 | Very High | High | Strong | Strong |
|  | 358 | 419 | Very High | High | Strong | Strong |
| **S-protein** | 281 | 337 | Very High | High | Strong | Strong |
|  | 533 | 617 | High | Medium | Strong | Strong |
|  | 925 | 1004 | High | High | Strong | None |
|  | 1145 | 1232 | Very High | High | Strong | Strong |

**Figure 4.**
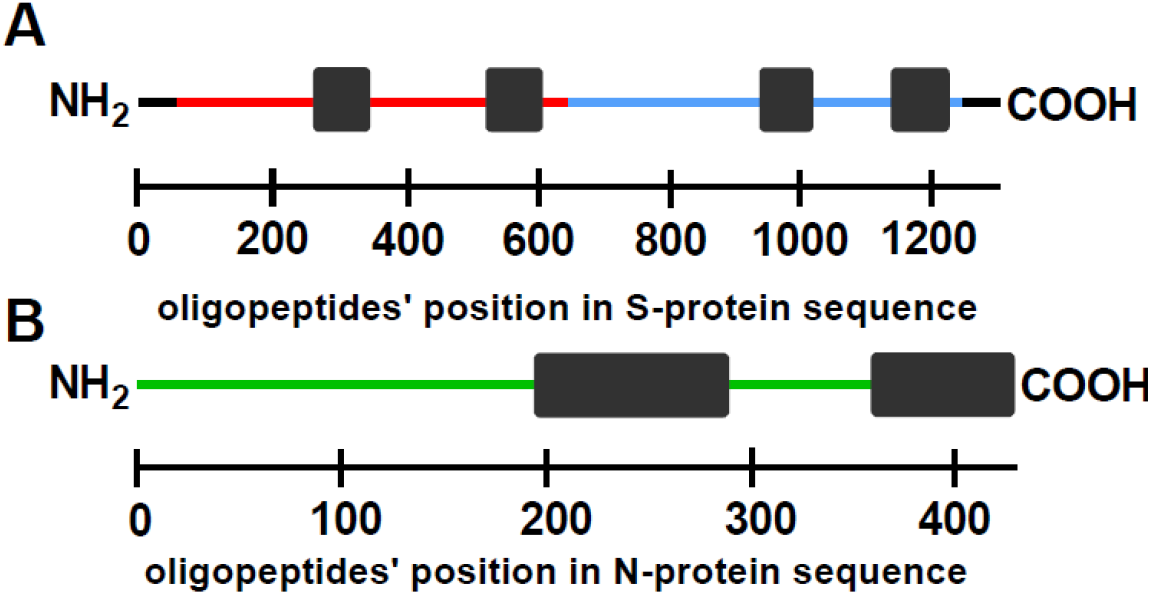
Representation of highly immunogenic regions with strong cross-reacting *IgG hot-spot* on sequences of proteins N and S. Representation of highly immunogenic regions with strong cross-reacting hot-spots (as black bars) on S protein (a) and N protein (b). Immunogenic regions efficiently induce specific IgG production in SARS-CoV-2 infected patients and these were identified by immunoprecipitation of the oligopeptide library representing Alpha and Delta (N_alpha_ = 15, N_delta_=8) SARS-CoV-2 variants (VirScan technology). Cross-reacting IgG hot spots are regions of protein frequently recognized in many variants by cross-reacting antibodies from patients. IgG hot-spots were identified by immunoprecipitation of the library containing SARS-CoV-2 variants’ oligopeptides (VirScan technology). Immunoprecipitation was conducted with sera from patients hospitalized due to SARS-CoV-2 infection (Alpha or Delta). Black boxes represent regions of respective protein with high or very high immunogenicity (p<0.05 and p<0.001, respectively) and strong cross-reacting hot-spots (p<0.001). P-values are calculated between experimental data and statistical binomial distribution model assuming random distribution. Red, blue and green lines represent S1, S2 subunits of S protein and N protein, respectively.

## Discussion

Potential significance of cross-reactivity that may characterize (or not) specific regions within SARS-CoV-2 proteins is linked to the practical application of these protein products. In N protein, cross-reactions increase its applicability as the diagnostic target in serological testing. For this reason, cross-reactivity hot-spots represent the regions where new mutations can probably be tolerated, without losing diagnostic applicability. In S protein, the major importance relates to anti-COVID-19 vaccines and to the protective potential of antibodies targeting this protein. Here, cross-reacting hot-spots represent regions where new mutations of the virus are relatively ‘safe’ for the human population; that is, the probability of immune evasion of a new variant with a mutation in a cross-reactivity hot-spot is lower. Regions outside the cross-reacting hot-spots, in turn, indicate higher probability of immune evasion by a new variant of that type.

Of note, this study has been focused on linear epitopes; thus the potential effect of mutations in structural epitopes could not be identified and it may still contribute to some specific effects of the immune response and protection. However, linear epitopes have been demonstrated as important targets in anti-SARS-CoV-2 protection ^5,15^. They are also of key importance for applications where a partial protein is used to elicit or detect the immune response. Therefore we propose the cross-reacting regions of N and S proteins in SARS-CoV-2 as the major targets in both diagnostics and vaccine design. Further, identified *IgG hot-spots* can be helpful in theoretical predictions of the potential for spread of newly identified variants, particularly when experimental data regarding a specific variant are not available yet. Variants with new mutations outside the cross-reacting hot-spots have the highest potential for immune evasion and eventually for spreading in the human population in spite of the vaccination rate or immunity acquired in infections caused by previous variants.

## Acknowledgements

This work was supported by the National Centre for Research and Development in Poland, grant no. SZPITALEJEDNOIMIENNE/48/2020. The authors are deeply grateful to all participants who agreed to act as serum donors in this study. Thank you for your help, support, understanding, and for your desire to contribute to scientific solutions to combat the disease.

No similar manuscript is or will be under consideration for publication elsewhere.

## Author Contributions

K.D. conceived the study. M.H. bioinformatic research preparation, K.D., K.G. designed and constructed the study groups. K.Ban., A.P., A.N., secured the day-to-day medical care and intervention for hospitalized patients at COVID-19 ward in Healthcare Centre Bolesławiec. K.G., I.R. conducted a research and investigation process, K.Bar, A.N., W.W. provided medical supervision of the project, K.D., K.G., K.Ban., A.P., A.N.,N.J., Z.K., T.K., M.K. contributed to the collection of data. M.H., analized, visualized and interpreted the results. M.H. and K.D. drafted the manuscript. K.D., W.W., critically revised the manuscript. All authors reviewed the manuscript and approved the final version.

## Data availability statement

The data underlying this study are not publicly available due to patient privacy issues. The data are available from the corresponding author upon reasonable request.

Supplementary information is available at Experimental&Molecular Medicine’s website.

Supplementary information accompanies the manuscript on the Experimental & Molecular Medicine’ website (http://www.nature.com/emm/).

## Supplementary material

**Supplementary Table S1.**
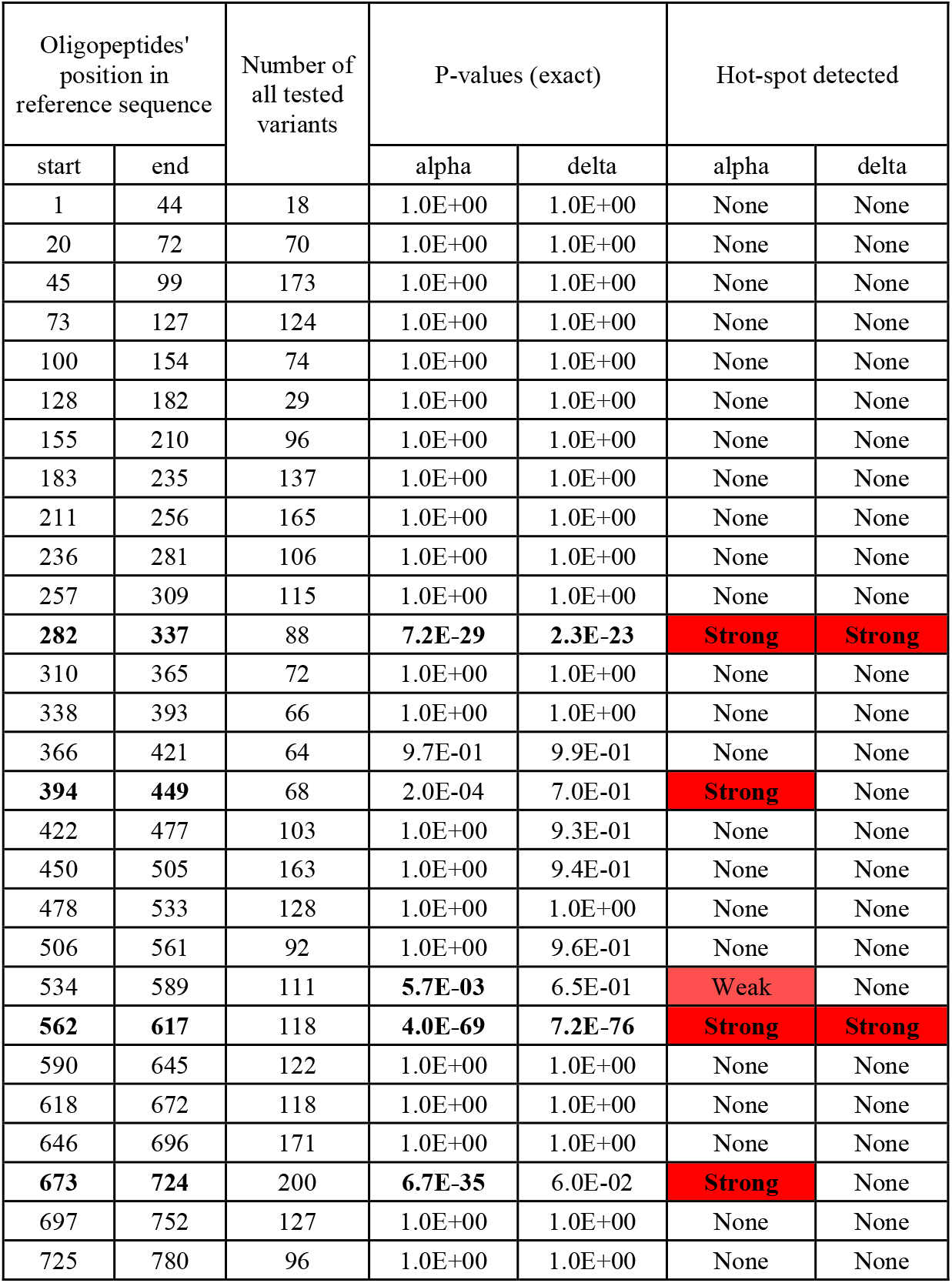
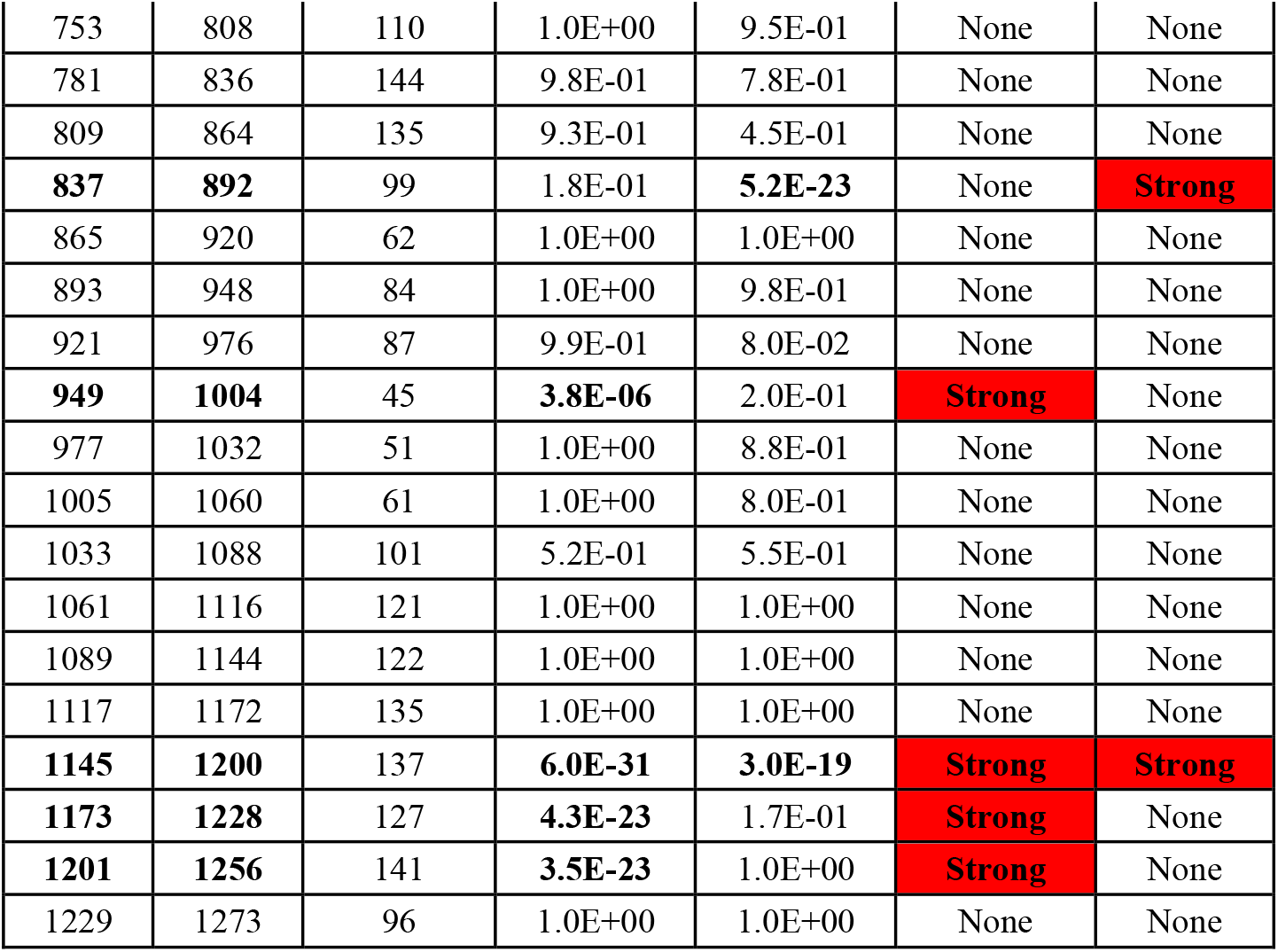
Cross-reacting IgG hot-spots in S protein of SARS-CoV-2. Cross-reacting IgG hot spots are regions of protein frequently recognized in many variants by cross-reacting antibodies. Cross-reactions of IgG antibodies were identified by immunoprecipitation of the library containing SARS-CoV-2 variants’ oligopeptides (VirScan technology). Immunoprecipitation was conducted with sera from patients hospitalized due to SARS-CoV-2 infection (Alpha or Delta). The fraction of cross-reacting variants of each oligopeptide was calculated from the total number of tested variants. Numbering consistent with the reference sequence (acc. no.: YP_009724390.1). P-values are calculated between measured data and the statistical binomial distribution model assuming random distribution of hot-spots in a protein.

**Supplementary Table S2.** Cross-reacting IgG hot-spots in N protein of SARS-CoV-2. Cross-reacting IgG hot spots are regions of protein frequently recognized in many variants by cross-reacting antibodies. Cross-reactions of IgG antibodies were identified by immunoprecipitation of the library containing SARS-CoV-2 variants’ oligopeptides (VirScan technology). Immunoprecipitation was conducted with sera from patients hospitalized due to SARS-CoV-2 infection (Alpha or Delta). The fraction of cross-reacting variants of each oligopeptide was calculated from the total number of tested variants. Numbering consistent with the reference sequence (acc. no.: YP_009724397.2). P-values are calculated between measured data and the statistical binomial distribution model assuming random distribution of hot-spots in a protein.

| Oligopeptides position in reference sequence |  | Number of all tested variants | P-values (exact) |  | Cross-reacting hot-spots |  |
| --- | --- | --- | --- | --- | --- | --- |
| start | end |  | alpha | delta | alpha | delta |
| 1 | 56 | 243 | 1.0E+00 | 1.0E+00 | None | None |
| 29 | 81 | 189 | 1.0E+00 | 9.3E-01 | None | None |
| **57** | **109** | 129 | 9.9E-01 | **5.4E-05** | None | **Strong** |
| 82 | 137 | 89 | 1.0E+00 | 1.0E+00 | None | None |
| 110 | 165 | 126 | 1.0E+00 | 1.0E+00 | None | None |
| **138** | **193** | 186 | **1.5E-09** | **8.1E-03** | **Strong** | Weak |
| **166** | **220** | 549 | **2.0E-41** | 1.0E+00 | **Strong** | None |
| 194 | 245 | 562 | 1.0E+00 | 1.0E+00 | None | None |
| **221** | **273** | 184 | **2.0E-28** | **5.3E-50** | **Strong** | **Strong** |
| 246 | 301 | 123 | 9.7E-01 | 9.5E-01 | None | None |
| 274 | 329 | 103 | 1.0E+00 | 1.0E+00 | None | None |
| 302 | 357 | 112 | 1.0E+00 | 1.0E+00 | None | None |
| 330 | 377 | 141 | 1.0E+00 | 1.0E+00 | None | None |
| **358** | **404** | 272 | **1.2E-04** | **1.0E-11** | **Strong** | **Strong** |
| **378** | **419** | 173 | **1.9E-06** | 8.7E-03 | **Strong** | Weak |

**Supplementary Table S3.** Immunogenicity and cross-reacting IgG hot-spots for M and E protein. Immunogenicity and cross-reactivity were detected by VirScan technology in sera of patients hospitalized due to alpha or delta variant SARS-CoV-2 infection. Below is presented the fraction of patients immunized to a region of reference SARS-CoV-2 protein that recognized the given region significantly more strongly. Also, we show a fraction of protein variants recognized by tested sera for each tested region of proteins. Red shows detected cross-reacting IgG hot spot – regions of protein that in multiple, natural proteins’ variants were efficiently bound by antibodies induced by tested sera.

| Protein name | Tested oligopeptides' position in ref. sequence [aa no.] | No. of variants tested | Fraction of patients immunized to a region [%] |  | Fraction of protein variants recognized by tested sera [%] |  |
| --- | --- | --- | --- | --- | --- | --- |
|  |  |  | alpha | delta | alpha | delta |
| M-protein | start |  |  |  |  |  |
|  | 1 | 65 | 26.7 | 50.0 | 6.2 | 4.6 |
|  | 29 | 100 | 13.3 | 0.0 | 21.0 | 3.0 |
|  | 57 | 85 | 0.0 | 0.0 | 5.9 | 0.0 |
|  | 85 | 77 | 0.0 | 0.0 | 0.0 | 0.0 |
|  | 113 | 27 | 26.7 | 50.0 | 0.0 | 0.0 |
|  | 143 | 78 | 0.0 | 0.0 | 1.3 | 0.0 |
|  | 171 | 61 | 20.0 | 25.0 | 0.0 | 0.0 |
|  | 199 | 61 | 0.0 | 0.0 | 4.9 | 1.6 |
| E-protein | 1 | 88 | 0.0 | 0.0 | 0.0 | 0.0 |
|  | 29 | 61 | 0.0 | 0.0 | 0.0 | 0.0 |
|  | 57 | 85 | 0.0 | 0.0 | 0.0 | 0.0 |

**Supplementary Table S4.**
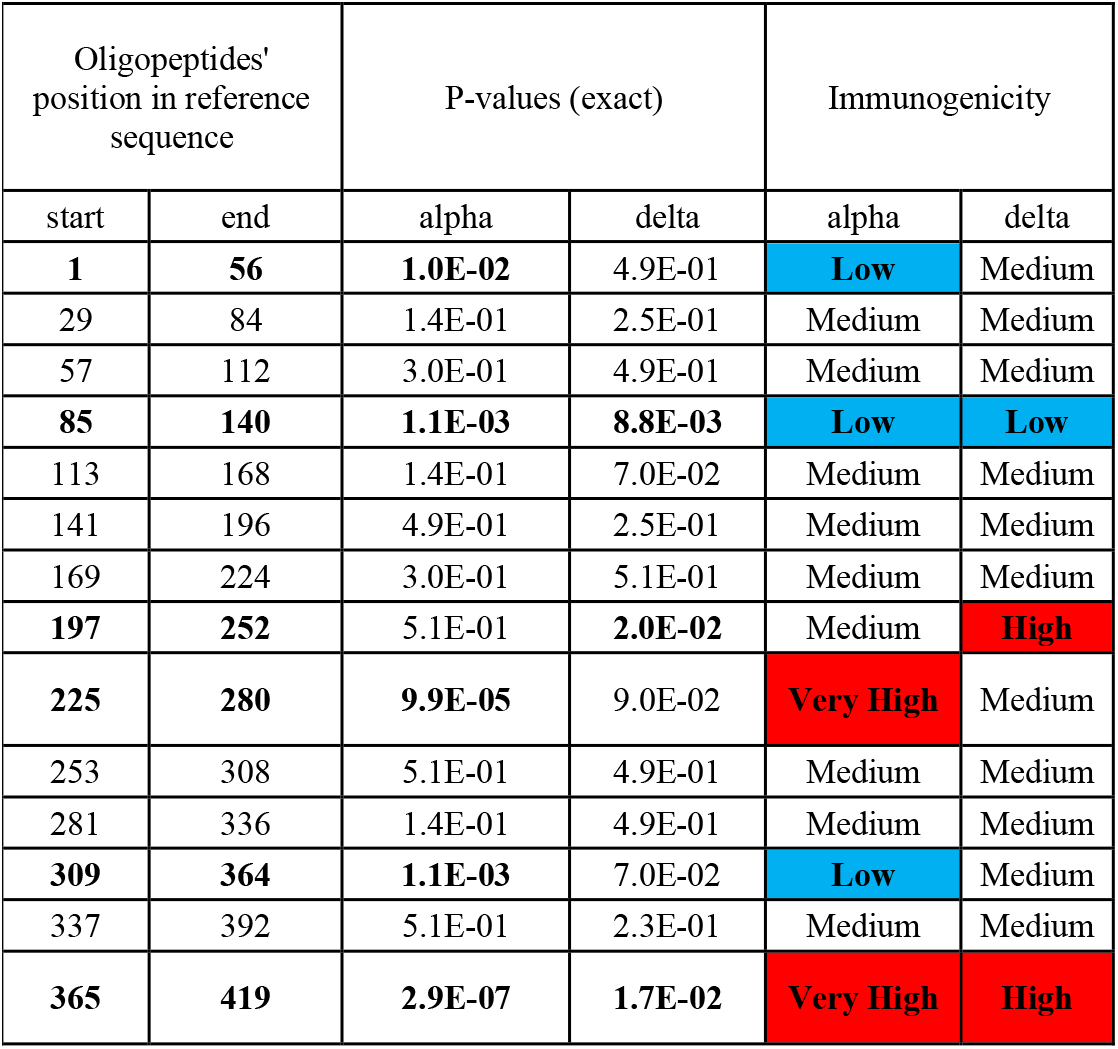
Immunogenic regions of SARS-CoV-2 protein N. Immunogenic regions efficiently induce specific IgG production in SARS-CoV-2 infected patients. Immunogenic regions were identified by immunoprecipitation of the oligopeptide library representing Alpha and Delta SARS-CoV-2 variants (VirScan technology). Immunoprecipitation was conducted with sera from patients hospitalized due to SARS-CoV-2 infection (Alpha or Delta). Red color represents regions of high and very high immunogenicity (p<0.05 and p<0.001, respectively). Blue color represents regions of low immunogenicity (p<0.05). P-value calculated between experimental data and statistical binomial distribution model assuming random distribution of hot-spot regions in proteins.

| Oligopeptides' position in reference sequence |  | P-values (exact) |  | Immunogenicity |  |
| --- | --- | --- | --- | --- | --- |
| start | end | alpha | delta | alpha | delta |
| **1** | **56** | **1.0E-02** | 4.9E-01 | **Low** | Medium |
| 29 | 84 | 1.4E-01 | 2.5E-01 | Medium | Medium |
| 57 | 112 | 3.0E-01 | 4.9E-01 | Medium | Medium |
| **85** | **140** | **1.1E-03** | **8.8E-03** | **Low** | **Low** |
| 113 | 168 | 1.4E-01 | 7.0E-02 | Medium | Medium |
| 141 | 196 | 4.9E-01 | 2.5E-01 | Medium | Medium |
| 169 | 224 | 3.0E-01 | 5.1E-01 | Medium | Medium |
| **197** | **252** | 5.1E-01 | **2.0E-02** | Medium | **High** |
| **225** | **280** | **9.9E-05** | 9.0E-02 | **Very High** | Medium |
| 253 | 308 | 5.1E-01 | 4.9E-01 | Medium | Medium |
| 281 | 336 | 1.4E-01 | 4.9E-01 | Medium | Medium |
| **309** | **364** | **1.1E-03** | 7.0E-02 | **Low** | Medium |
| 337 | 392 | 5.1E-01 | 2.3E-01 | Medium | Medium |
| **365** | **419** | **2.9E-07** | **1.7E-02** | **Very High** | **High** |

**Supplementary Table S5:** Immunogenic regions in protein S. Immunogenic regions efficiently induce specific IgG production in SARS-CoV-2 infected patients. Immunogenic regions were identified by immunoprecipitation of the oligopeptide library representing Alpha and Delta SARS-CoV-2 variants (VirScan technology). Immunoprecipitation was conducted with sera from patients hospitalized due to SARS-CoV-2 infection (Alpha or Delta). Red color represents regions of high and very high immunogenicity (p<0.05 and p<0.001, respectively). Blue color represents regions of low immunogenicity (p<0.05). P-value calculated between experimental data and statistical binomial distribution model assuming random distribution of hot-spot regions in proteins.

| Oligopeptides' position in reference sequence |  | P-values (exact) |  | Immunogenicity |  |
| --- | --- | --- | --- | --- | --- |
| start | end | alpha | delta | alpha | delta |
| **1** | **56** | **0.0E+00** | **5.0E-02** | **Low** | **Low** |
| **29** | **84** | **3.0E-02** | **5.0E-02** | **Low** | **Low** |
| **57** | **112** | 1.0E-01 | **5.0E-02** | Medium | **Low** |
| **85** | **140** | **3.0E-02** | **5.0E-02** | **Low** | **Low** |
| **113** | **168** | **0.0E+00** | **5.0E-02** | **Low** | **Low** |
| **141** | **196** | **0.0E+00** | **5.0E-02** | **Low** | **Low** |
| **169** | **224** | 4.7E-01 | **1.6E-02** | Medium | **High** |
| 197 | 252 | 5.3E-01 | 2.2E-01 | Medium | Medium |
| **225** | **280** | **0.0E+00** | 2.2E-01 | **Low** | Medium |
| 253 | 308 | 1.0E-01 | 2.2E-01 | Medium | Medium |
| **281** | **336** | **1.5E-05** | **1.6E-02** | **Very High** | **High** |
| 309 | 364 | 4.7E-01 | 2.2E-01 | Medium | Medium |
| 337 | 392 | 2.6E-01 | 2.2E-01 | Medium | Medium |
| 365 | 420 | 5.3E-01 | 5.0E-01 | Medium | Medium |
| 393 | 448 | 1.6E-01 | 5.0E-01 | Medium | Medium |
| 421 | 476 | 2.6E-01 | 5.0E-01 | Medium | Medium |
| **449** | **504** | **3.0E-02** | 5.0E-01 | **Low** | Medium |
| **477** | **532** | **3.0E-02** | 2.2E-01 | **Low** | Medium |
| 505 | 560 | 1.6E-01 | 5.0E-01 | Medium | Medium |
| **533** | **588** | **5.3E-03** | 5.0E-01 | **High** | Medium |
| **561** | **616** | **1.0E-03** | 7.0E-02 | **High** | Medium |
| **589** | **644** | **3.0E-02** | 2.2E-01 | **Low** | Medium |
| 617 | 672 | 1.0E-01 | 5.0E-01 | Medium | Medium |
| 645 | 700 | 1.0E-01 | 2.2E-01 | Medium | Medium |
| 673 | 728 | 2.0E-02 | 2.3E-01 | Medium | Medium |
| **701** | **756** | **0.0E+00** | 7.0E-02 | **Very High** | Medium |
| 729 | 784 | 2.6E-01 | 2.2E-01 | Medium | Medium |
| **757** | **812** | 6.0E-02 | **1.6E-02** | Medium | **High** |
| **785** | **840** | **1.5E-04** | **2.0E-03** | **Very High** | **High** |
| **813** | **868** | **1.0E-03** | **2.0E-03** | **High** | **High** |
| 841 | 896 | 5.3E-01 | 5.0E-01 | Medium | Medium |
| **869** | **924** | **0.0E+00** | **5.0E-02** | **Low** | **Low** |
| 897 | 952 | 5.3E-01 | 2.3E-01 | Medium | Medium |
| **925** | **980** | **5.3E-03** | **1.6E-02** | **High** | **High** |
| 953 | 1008 | 5.3E-01 | 2.3E-01 | Medium | Medium |
| 981 | 1036 | 2.6E-01 | 2.3E-01 | Medium | Medium |
| **1009** | **1064** | **0.0E+00** | 5.0E-01 | **Low** | Medium |
| 1037 | 1092 | 4.7E-01 | 5.0E-01 | Medium | Medium |
| 1065 | 1120 | 6.0E-02 | 5.0E-01 | Medium | Medium |
| 1093 | 1148 | 1.0E-01 | 2.2E-01 | Medium | Medium |
| **1121** | **1176** | 1.0E-01 | **5.0E-02** | Medium | **Low** |
| **1149** | **1204** | **9.6E-07** | **2.0E-03** | **Very High** | **High** |
| **1177** | **1232** | **1.5E-05** | **2.0E-03** | **Very High** | **High** |
| **1205** | **1260** | **3.0E-02** | 2.2E-01 | **Low** | Medium |
| **1233** | **1273** | **0.0E+00** | **5.0E-02** | **Low** | **Low** |

**Figure S1.**
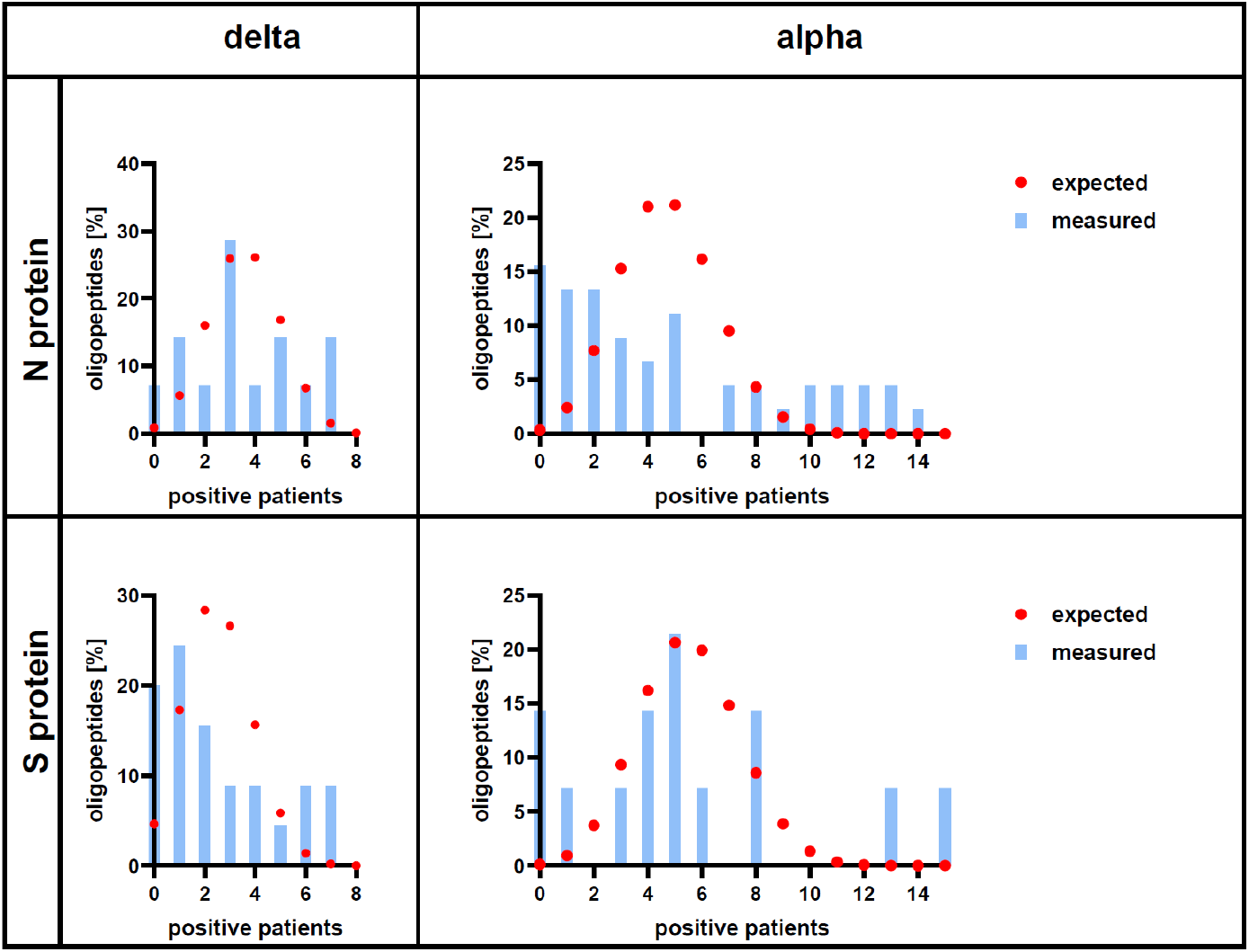
Comparison of binomial distribution model and experimental results of testing proteins’ S&N immunogenicity. X-axis shows the number of patients in a tested group (hospitalized for Alpha or Delta variant) whose sera bind the same oligopeptide. Y-axis shows the expected chance of a given number of patients testing positive for the same, given oligopeptide. Blue bars represent measured, experimental frequencies of the given number of positive patients. Red dots represent the chance that measuring this specific number of patients testing positive would occur randomly (binomial distribution model).

## References

1. Jacobs, J. L., Haidar, G. & Mellors, J. W. COVID-19: Challenges of Viral Variants. Annu. Rev. Med. 74, (2022).

2. Akkız, H. The Biological Functions and Clinical Significance of SARS-CoV-2 Variants of Corcern. Front. Med. 9, (2022).

3. Yewdell, J. W. Antigenic drift: Understanding COVID-19. Immunity 54, 2681–2687 (2021).

4. Yang, H. & Rao, Z. Structural biology of SARS-CoV-2 and implications for therapeutic development. Nat. Rev. Microbiol. 2021 1911 19, 685–700 (2021).

5. Tretyn, A. et al. Differences in the Concentration of Anti-SARS-CoV-2 IgG Antibodies Post-COVID-19 Recovery or Post-Vaccination. Cells 10, (2021).

6. Zhang, Y. et al. Diagnostic Value of Nucleocapsid Protein in Blood for SARS-CoV-2 Infection. Clin. Chem. 68, 240–248 (2021).

7. Xu, G. J. et al. Comprehensive serological profiling of human populations using a synthetic human virome. Science (80-.). 348, (2015).

8. Sievers, F. et al. Fast, scalable generation of high-quality protein multiple sequence alignments using Clustal Omega. Mol. Syst. Biol. 7, 539 (2011).

9. Goujon, M. et al. A new bioinformatics analysis tools framework at EMBL–EBI. Nucleic Acids Res. 38, W695–W699 (2010).

10. McWilliam, H. et al. Analysis Tool Web Services from the EMBL-EBI. Nucleic Acids Res. 41, W597–W600 (2013).

11. Harhala, M., Gembara, K., Nelson, D. C., Miernikiewicz, P. & Dabrowska, K. Immunogenicity of Endolysin PlyC. 1–12 (2022).

12. Langmead, B. & Salzberg, S. L. Fast gapped-read alignment with Bowtie 2. Nat. Methods 9, 357–359 (2012).

13. Langmead, B., Trapnell, C., Pop, M. & Salzberg, S. L. Ultrafast and memory-efficient alignment of short DNA sequences to the human genome. Genome Biol. 10, (2009).

14. Elshabrawy, H. A., Coughlin, M. M., Baker, S. C. & Prabhakar, B. S. Human Monoclonal Antibodies against Highly Conserved HR1 and HR2 Domains of the SARS-CoV Spike Protein Are More Broadly Neutralizing. PLoS One 7, e50366 (2012).

15. Gao, W. et al. Characterization and analysis of linear epitopes corresponding to SARS-CoV-2 outbreak in Jilin Province, China. J. Med. Virol. 95, (2023).

